## Supplementary material for "Rotavirus A Genome Segments Show Distinct Segregation and Codon Usage Patterns": SI figures 1 to 11: SI_Trees1to5.docx

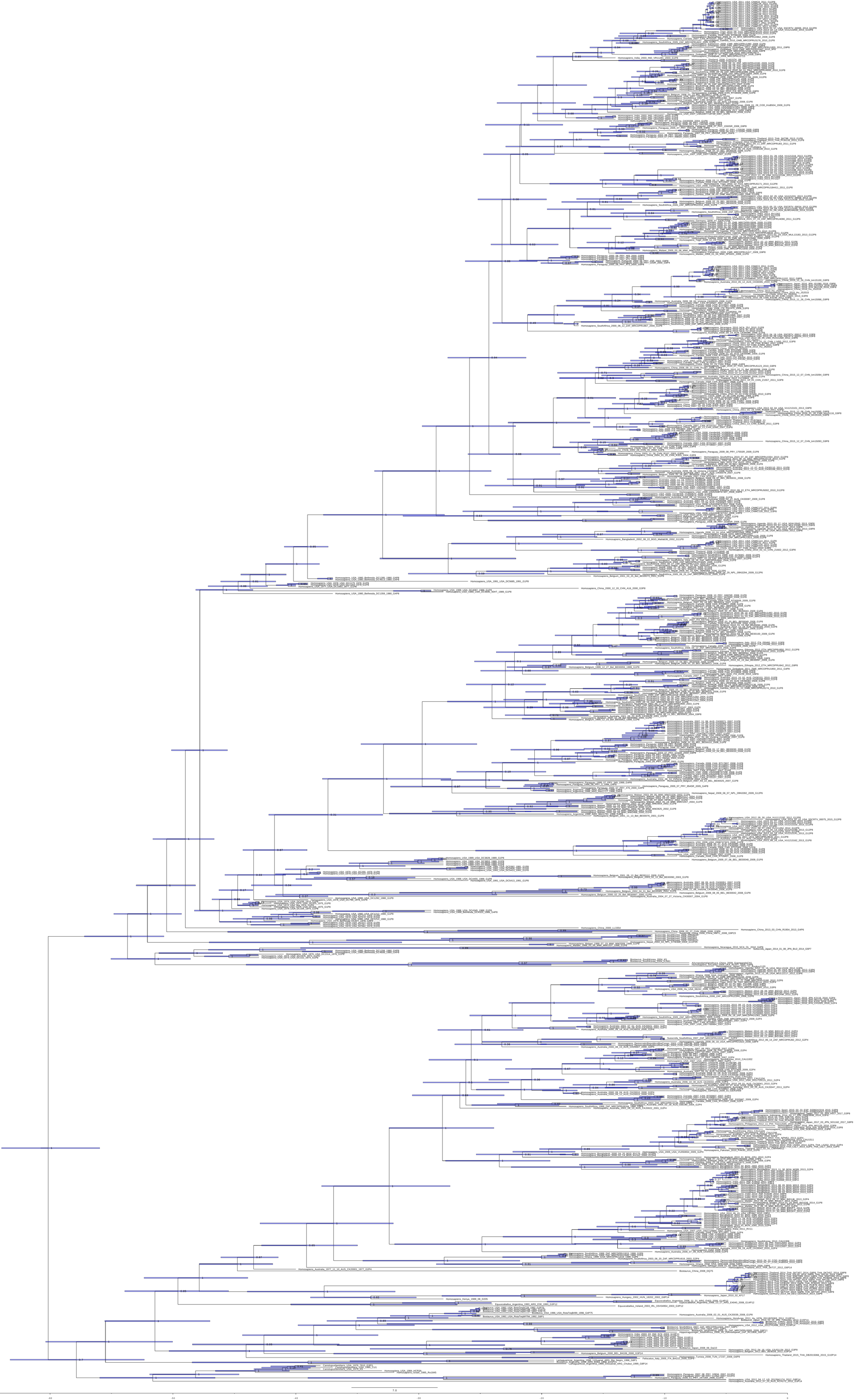

SI Figure 1. Time-scaled Phylogeny of Segment 1. Time-scaled axis is in years (since 2017). Node divergence date confidence intervals are shown by node bars. Posterior probabilities are labeled on the nodes. Strain names are contained in the tip labels.

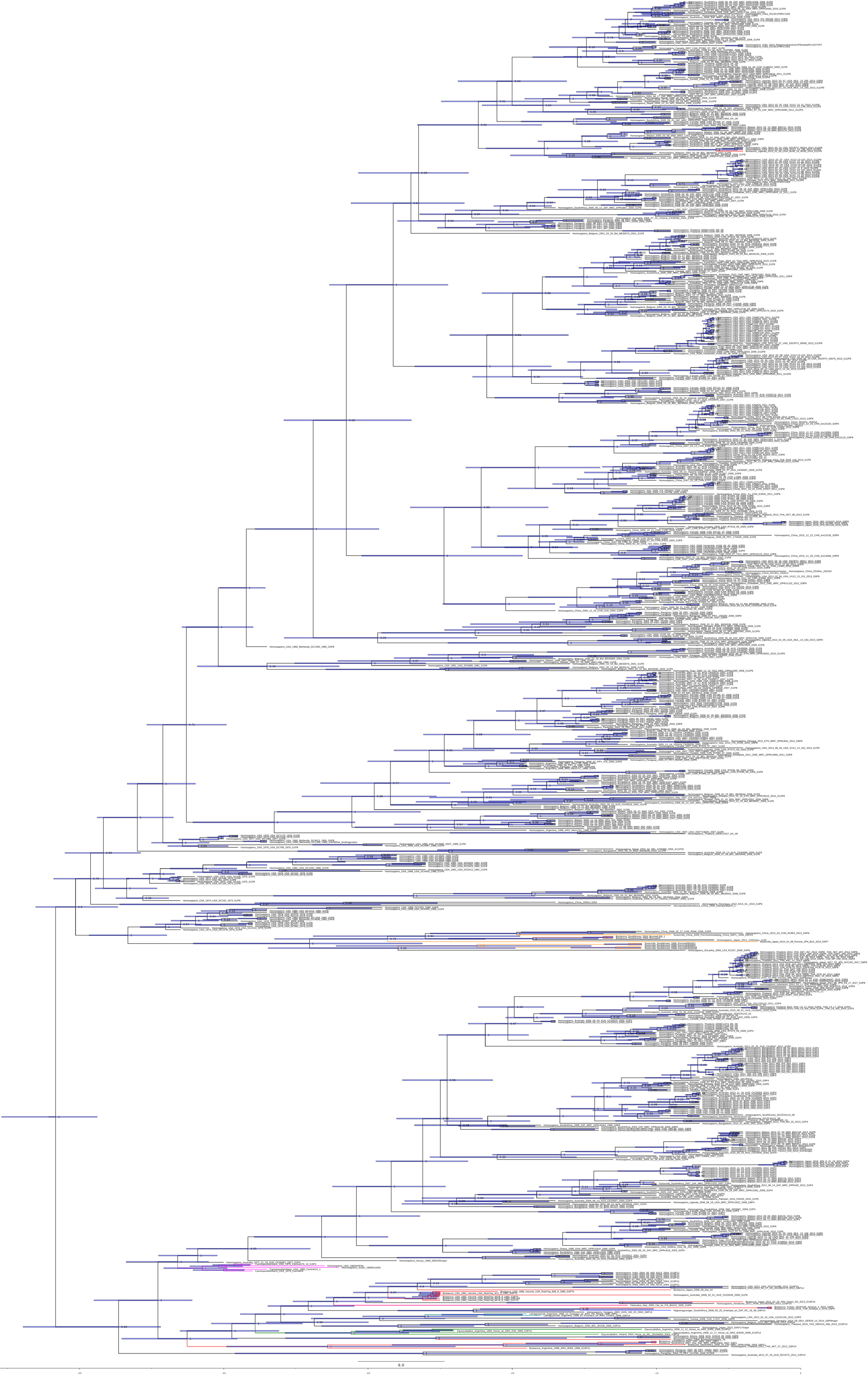

SI Figure 2. Time-scaled Phylogeny of Segment 2. Time-scaled axis is in years (since 2017). Node divergence date confidence intervals are shown by node bars. Posterior probabilities are labeled on the nodes. Strain names are contained in the tip labels.

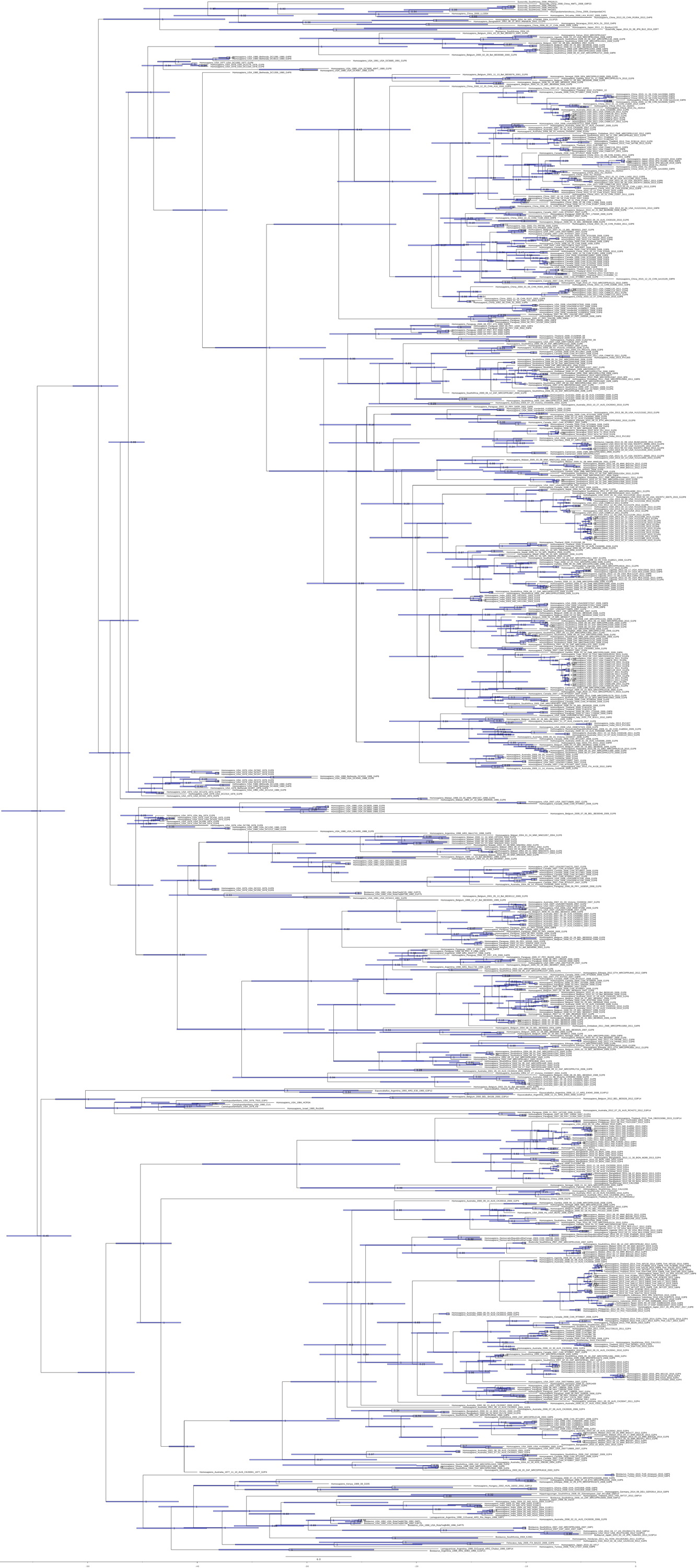

SI Figure 3. Time-scaled Phylogeny of Segment 3. Time-scaled axis is in years (since 2017). Node divergence date confidence intervals are shown by node bars. Posterior probabilities are labeled on the nodes. Strain names are contained in the tip labels.

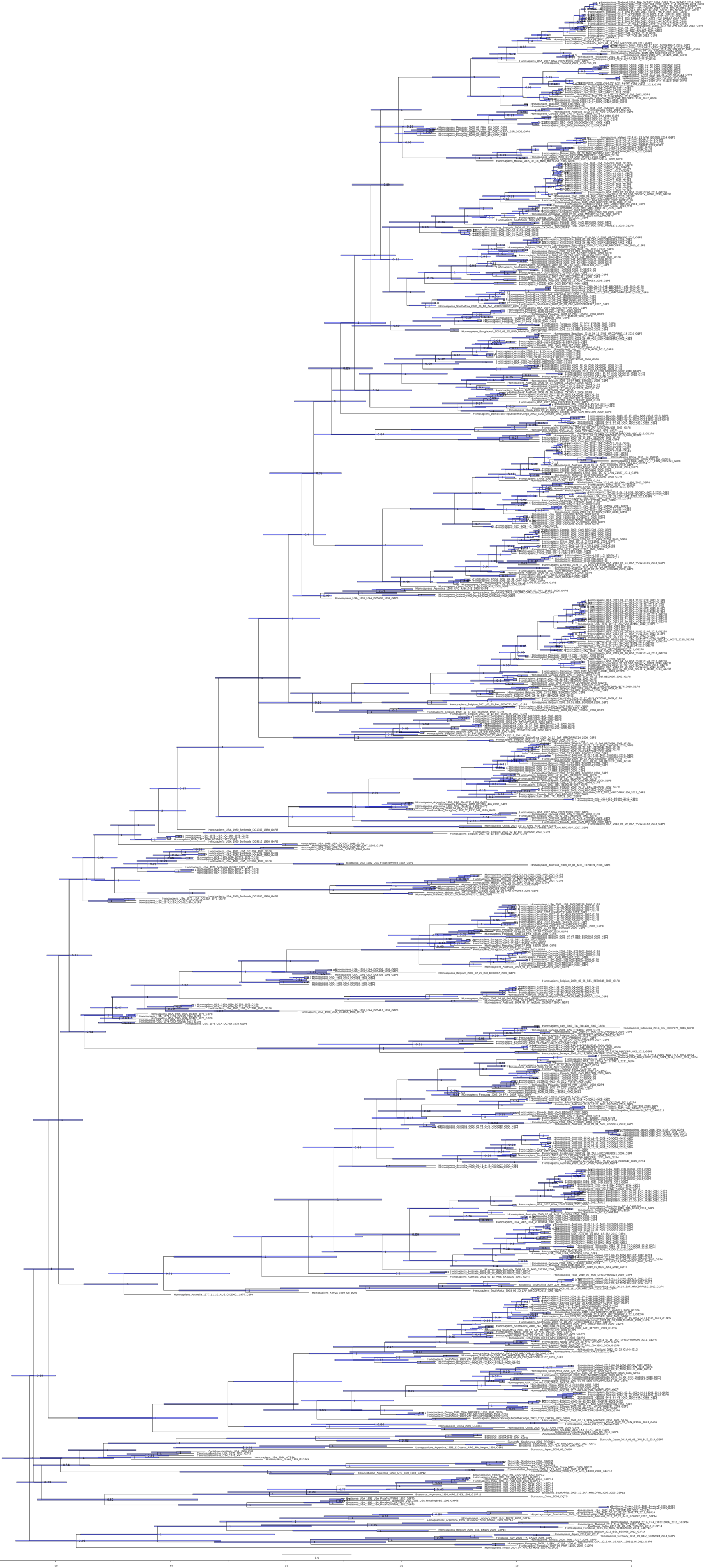

SI Figure 4. Time-scaled Phylogeny of Segment 4. Time-scaled axis is in years (since 2017). Node divergence date confidence intervals are shown by node bars. Posterior probabilities are labeled on the nodes. Strain names are contained in the tip labels.

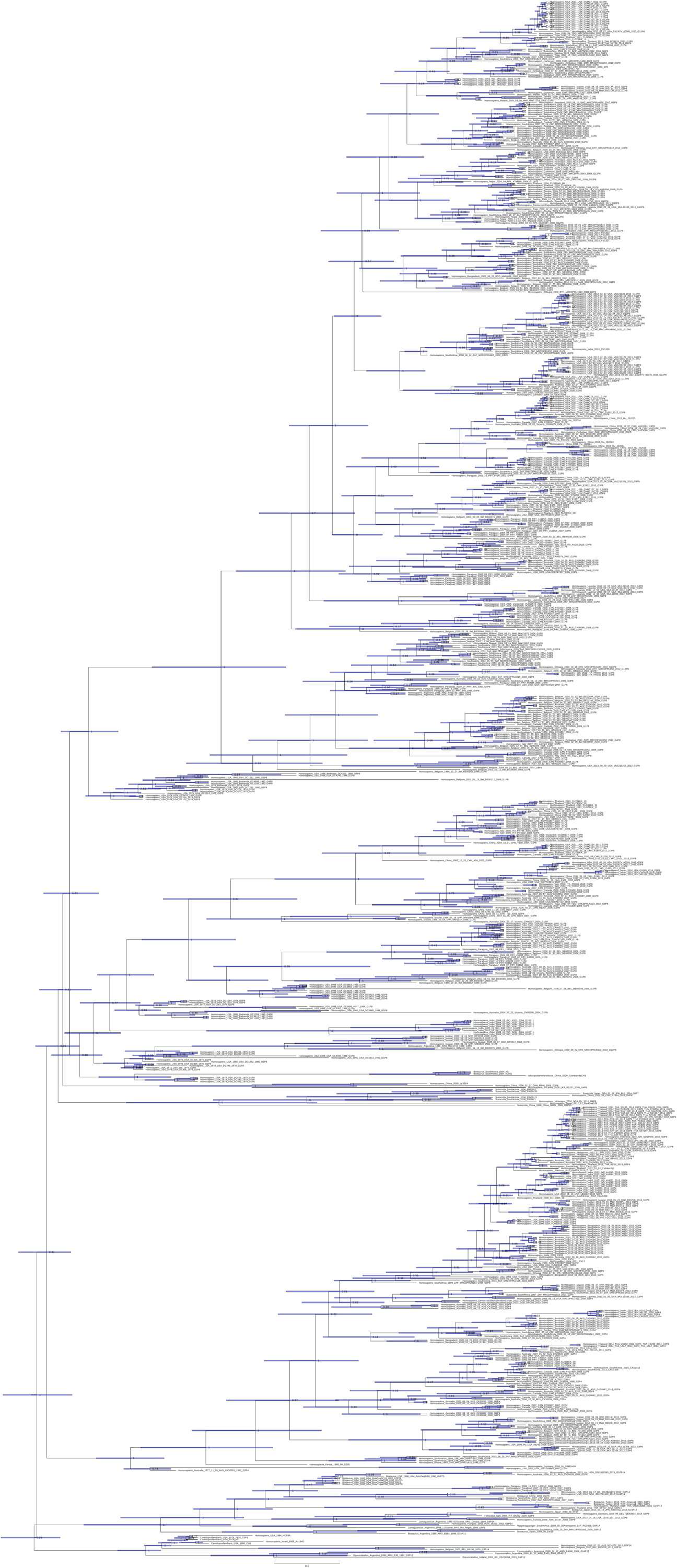

SI Figure 5. Time-scaled Phylogeny of Segment 5. Time-scaled axis is in years (since 2017). Node divergence date confidence intervals are shown by node bars. Posterior probabilities are labeled on the nodes. Strain names are contained in the tip labels.
